# Polymorphisms in human APOBEC3H differentially regulate ubiquitination and antiviral activity

**DOI:** 10.1101/2020.02.07.939439

**Authors:** Nicholas M. Chesarino, Michael Emerman

## Abstract

The APOBEC3 family of cytidine deaminases are an important part of the host innate immune defense against endogenous retroelements and retroviruses like human immunodeficiency virus (HIV). *APOBEC3H* (*A3H*) is the most polymorphic of the human *APOBEC3* genes, with four major haplotypes circulating in the population. Haplotype II is the only antivirally-active variant of *A3H*, while the majority of the population possess independently destabilizing polymorphisms present in haplotype I (R105G) and haplotypes III and IV (N15del). Here, we show that instability introduced by either polymorphism is positively correlated with degradative ubiquitination, while haplotype II is protected from this modification. Inhibiting ubiquitination by mutating all of the A3H lysines increased expression of haplotypes III and IV, but these stabilized forms of haplotype III and IV had a strict nuclear localization, and did not incorporate into virions, nor exhibit antiviral activity, thus separating stabilization from function. On the other hand, the instability and functional deficiencies of haplotype III could be rescued by fusion to haplotype II, supporting a model by which antiviral A3H is actively stabilized through a cytoplasmic retention mechanism. Thus, the evolutionary loss of A3H activity in many humans involves functional deficiencies independent of protein stability.

## 1. Introduction

Restriction factors are a critical component of the innate immune response to cellular infection by viruses by inhibiting various stages of the viral replication cycle. The APOBEC3 (A3) family of cytidine deaminases are potent restriction factors against endogenous retroelements and retroviruses, with four members of this family (A3D, A3F, A3G, and A3H) demonstrating considerable antiviral activity against lentiviruses such as Simian Immunodeficiency Virus (SIV) and Human Immunodeficiency Virus (HIV) [1]. The antiviral A3 proteins incorporate into nascent virions and act during the reverse transcription stage of retroviral infection to convert cytosines to uracils in viral single-stranded DNA [2]. The result of successful A3 activity is lethal hypermutation of the retroviral genome, preventing the establishment of a productive infection. Thus, the A3 proteins act in concert to provide a barrier to both cross-species transmission and within-host cell-to-cell spread of infection.

All of the *A3* genes display strong signatures of positive selection throughout primate evolution, suggesting a longstanding genetic conflict between this family of restriction factors and the pathogens and endogenous retroelements they restrict [3–5]. To overcome A3 restriction, lentiviruses have evolved the accessory protein, Vif, that potently inhibits the function of all antiviral A3 proteins simultaneously [6,7]. Vif hijacks a host cellular E3 ubiquitin ligase complex and acts as a substrate receptor for A3 proteins, promoting their ubiquitination and subsequent degradation via the proteasome [8–11]. The direct antagonistic relationship between host A3 and lentiviral Vif underlies an evolutionary arms race that selects for mutations in A3 that can escape Vif antagonism, and mutations in Vif that regain A3 binding [12,13]. Therefore, the A3 proteins confer selective pressure on lentiviruses to evolve Vif proteins that can overcome the host A3 defense.

Despite the selective pressure to retain host defense systems like the *A3* genes, restriction factors can become less effective due to circulating polymorphisms that reduce inherent antiviral function. It is possible that such polymorphisms may arise and become fixed in the population upon the loss of a selective pathogenic pressure, especially if maintenance of a particular antiviral gene is deleterious to the host. A3H is a notable example of such a polymorphic restriction factor in humans, with four major *A3H* variants (haplotypes I, II, III, and IV) circulating in the human population, and at least eight other minor haplotypes identified to date [14]. Among these, haplotype II is the only variant to encode a protein with potent activity against lentiviruses [15–19]. In contrast, the protein encoded by haplotype I is far less antiviral and stable, and proteins encoded by haplotypes III and IV are not detectable at all [15–18,20,21]. Evolutionary evidence suggests that human A3H lost activity in two independent events since the last common ancestor with chimpanzees [16]. Thus, the destabilization of haplotypes I, III, and IV are a consequence of two independent single nucleotide polymorphisms, R105G in haplotype I and a deletion of codon 15, N15del, in haplotypes III and IV. Each mutation is sufficient to render A3H ineffective against HIV [15,16,18]. Importantly, the differences in protein expression are not explained by differences in transcript levels, as mRNA levels are comparable for both overexpressed stable and unstable alleles [22] as well as endogenous transcripts of various alleles measured in primary T lymphocytes [17]. Rather, expression differences are due to reduced protein half-lives [16].

Recent structural studies of pig-tailed macaque [23], human [24,25], and chimpanzee [26] A3H have established a unique RNA interaction mechanism central to the antiviral function of A3H. Two A3H monomers interact with opposite sides of a short RNA duplex, and the A3H monomers in this complex interact primarily with the RNA duplex and not with one another. Importantly, mutagenesis of residues in the interaction interface between A3H and duplex RNA results in a decrease in A3H expression similar to the unstable human haplotypes [25–27]. Thus, formation of the A3H-RNA complex may be a critical regulator of A3H stability. However, the exact mechanism by which R105G destabilizes A3H is not well understood, and even less is known of the effect of the N15del mutation in haplotypes III and IV.

In order to better understand the processes underlying the differential expression of different A3H haplotypes, we asked how ubiquitination, a posttranslational modification most widely known for its role in protein degradation, differs between A3H haplotypes. We found that the rates of ubiquitination were greater in unstable haplotypes I, III, and IV, while the stable haplotype II was largely protected from this modification. By genetically inhibiting ubiquitination through lysine mutagenesis, we were able to recover expression of the N15del-containing haplotypes III and IV to levels comparable to haplotype I. However, despite increasing protein expression, these changes did not restore antiviral activity of any of these haplotypes against HIV-1. Rather, stabilized versions of the proteins encoded by haplotypes III and IV were strictly localized to the nucleus and were unable to package into budding virions. Thus, the R105G and N15del mutations in A3H result in functional defects that cannot be restored by inhibiting ubiquitination alone. On the other hand, by fusing the stable and antiviral haplotype II to haplotype III, the deficiencies of the N15del mutation on expression, localization, and antiviral activity were reversed. Taken together, these results suggest that both A3H stability and activity are tightly regulated by loss of function mutations, and selection for these mutations may serve to protect the host in the absence of a pathogenic pressure.

## 2. Materials and Methods

### 2.1. Plasmids

Plasmids containing A3H haplotypes I through IV were previously described [16], and used as templates to generate C-terminally HA-tagged constructs. All constructs used in this study were cloned into pcDNA3.1 (Thermo Fisher) using EcoRI/XhoI restriction sites. Mutations were introduced using standard PCR or using the QuikChange Lightning Multi Site-Directed Mutagenesis Kit (Agilent, 210515). Double A3H fusion constructs I-I, II-II, I-II, and II-I were previously described [20]. The double A3H fusion constructs unique to this study, III-III, III-II, and II-III were made through amplification of haplotypes II and III with a 5’ or 3’ primer containing the linker sequence GGT GGT GGT GGT GGC GCC (Gly-Gly-Gly-Gly-Gly-Ala). 5’ and 3’ haplotype domains were digested using the KasI restriction site in the linker, and EcoRI (for 5’ domains) or XhoI (for 3’ domains). 5’ domains, 3’ domains, and the EcoRI/XhoI double digested pcDNA3.1 vector were joined using T4 DNA Ligase (New England BioLabs, M0202). Infectivity experiments were performed using the HIV-1 molecular clone pLAIΔenvLuc2Δvif, which has been previously described [28].

### 2.2. Cell lines and transfections

293T, HeLa, and SupT1 cells were obtained from ATCC (CRL-3216, CCL-2, and CRL-1942, respectively). 293T and HeLa cells were cultured in DMEM, high glucose (Gibco, 11965092) supplemented with 10% HyClone Bovine Growth Serum (GE Healthcare Life Sciences, SH30541.03) and 1X Penicillin-Streptomycin (Gibco, 15140122). SupT1 cells were cultured in RPMI-1640 (Gibco, 11875093) supplemented with 10% HyClone Fetal Bovine Serum (GE Healthcare, SH3091003) and 1X Penicillin-Streptomycin. Cell lines were mycoplasma free, as determined by the Fred Hutchinson Cancer Research Center Specimen Processing/Research Cell Bank core facility. Cells were incubated at 37°C and 5% CO_2_ in a humidified incubator and maintained for under thirty passages before returning to a lower passage stock.

Transfections were performed using *Trans*IT-LT1 Transfection Reagent (Mirus, MIR 2305) according to the manufacturer’s protocol. For western blotting, 293T cells were plated at 1.5×10^5^/mL 24 h prior to transfection. Transfections in 12-well plates were performed using 0.5 μg/well of A3H plasmid and 3 μL/well of transfection reagent. For ubiquitination experiments, 6-well plates were transfected with 1.0 μg/well of A3H plasmid, 1.0 μg/well of myc-Ubiquitin, and 6 μL/well of transfection reagent. For infectivity experiments, 293T cells in 96-well plates (3.75×10^4^/well) were reverse-transfected with 60 ng/well pLAIΔenvLuc2Δvif, 30 ng/well A3H plasmid, and 10 ng/well L-VSV-G. For immunofluorescence microscopy, cells were plated at 5.0×10^4^/mL on glass cover slips in a 12-well plate 24 h prior to transfection with 0.5 μg/mL of A3H plasmid and 3 μL/well of transfection reagent. In all cases, transfected cells were incubated for 48 h prior to downstream applications.

### 2.3. Viral infectivity assays

Viruses were propagated in 293T cells reverse-transfected with pLAIΔenvLuc2Δvif, A3H plasmid, and L-VSV-G for pseudotyping. Virus-containing supernatants were harvested 48 h after transfection, transferred to a V-bottom plate, and clarified of cells and debris by centrifugation at 1,000xG for 3 min at 25°C. 10 μL of supernatant was added to flat-bottom 96-well plates containing SupT1 cells (3.75×10^4^ and 90 μL/well) pretreated with 20 μg/mL DEAE/Dextran and mixed by repipetting. 5 μL of supernatant was saved for quantification of reverse transcriptase (RT) activity, as described previously [29]. Infected SupT1 cells were incubated for 48 h and lysed in 100 μL of Bright-Glo Luciferase Reagent (Promega, E2610). Infection was assessed by luciferase activity using a LUMIstar Omega microplate luminometer (BMG Labtech), and raw luciferase values were normalized to 2,000 mU RT activity.

### 2.4. Immunoprecipitation and western blotting

For experiments involving MG132, cells were treated with 10 μM MG132 (Selleck Chem, S2619), or an equal volume of DMSO, in fresh media for 18 h after the initial 48 h transfection. For all other experiments, cells were harvested 48 h post-transfection. Cells were washed twice with PBS, and lysed on ice with RIPA buffer (50 mM Tris-HCl, pH 7.4, 150 mM NaCl, 1.0% Triton X-100, 0.5% sodium deoxycholate, 1 mM EDTA, 1 mM MgCl_2_) for 20 min. For ubiquitination assays, transfected cells were and lysed in 150 μL of 1% SDS buffer (50 mM Tris-HCl pH 7.4, 1.0% SDS) at 95°C for 10 min. Lysate was passed through a 30-gauge needle to sheer DNA. 50 μg of lysate was saved for input. An equal concentration of lysate for immunoprecipitation was diluted in 100 μL of 1% SDS buffer, which was further diluted in 900 μL ice cold RIPA buffer lacking SDS containing 15 μL EZview Red Anti-HA Affinity Gel (Sigma, E6779). Lysate was immunoprecipitated for 1 h at 4°C with nutation, washed three times with ice cold RIPA buffer lacking SDS, and eluted in 40 μL 2X Laemmli Sample Buffer (BIO-RAD, 1610737). Lysis buffers were supplemented with cOmplete Protease Inhibitor Cocktail (Roche, 11697498001), 10 μM MG132, and 50 μM PR-619 deubiquitinase inhibitor (Selleck Chem, S7130).

10 μg per lysate or 10 μL of immunoprecipitated sample was resolved on a NuPAGE 4-12% Bis-Tris Protein Gel (Invitrogen, NP0336). Western blotting was performed using the primary antibodies anti-HA (Proteintech, 51064-2-AP), anti-myc (Proteintech, 16286-1-AP), anti-A3H (P1H6-1, [28]), and anti-Lamin B1 (Proteintech, 66095-1-Ig) at a dilution of 1:2,000. Secondary antibodies StarBright Blue 520 Goat Anti-Rabbit IgG (BIO-RAD, 12005869) and StarBright Blue 700 Goat Anti-Mouse IgG (BIO-RAD, 12005866) were used at a dilution of 1:10,000. Densitometric analysis was performed using ImageJ software [30]. All immunoblots are representative of at least three independent experiments.

### 2.5. Fluorescence microscopy

Transfected cells on glass coverslips were washed twice with PBS and fixed in 4% paraformaldehyde, permeabilized in 0.1% Triton X-100 in PBS, and blocked in 2% Bovine Serum Albumin in PBS for 10 min each at room temperature. Primary antibodies anti-HA.11 (Clontech, 631207) or anti-A3H were used at a dilution at 1:500 and 1:50, respectively, in 0.1% Triton X-100 in PBS and incubated with cells for 20 min at room temperature. Cells were washed five times with 0.1% Triton X-100 in PBS prior to incubation with Alexa Fluor 488-labeled anti-mouse secondary antibody at a 1:1000 dilution in 0.1% Triton X-100 in PBS. Cells were again washed five times prior to mounting glass coverslips in ProLong Gold antifade reagent containing DAPI (Life Technologies, P36935). Images were obtained using a Nikon E800 microscope at 40X magnification.

## 3. Results

### 3.1. Rates of ubiquitination differ among A3H haplotypes

The majority of the human population possess A3H haplotypes that make unstable proteins with no antiviral activity. Haplotype I, containing the R105G mutation, has a global allele frequency of 46.4%, and haplotypes III and IV, containing N15del, make up a combined 30.2% of the population [14]. The protein encoded by A3H haplotype II has more steady-state expression than the protein encoded by haplotype I, and the proteins encoded by haplotypes III and IV are barely, if at all, detectable [15–18,31]. The question of why unstable and inactive variants of potent restriction factors occur at such high frequency in the human population and whether these inactive antiviral proteins could be manipulated to restore an innate immune function against HIV-1, prompted us to ask if A3H activity could be induced through stabilization, or if antiviral activity was lost independent of stability.

Previous work has shown that disruption of the proteasome with the drug MG132 modestly increases expression of both of the proteins encoded by A3H haplotypes I and II, suggesting that stability might be linked to proteasomal turnover [22,31]. We reexamined this question by evaluating the effect of MG132 on A3H haplotypes III and IV, which encode for the most poorly expressed proteins and contain the N15del polymorphism. We overexpressed each of the four HA-tagged A3H (A3H-HA) haplotypes in 293T cells, followed by treatment with the proteasomal inhibitor MG132 or a DMSO control. MG132 only slightly increased expression of haplotype II (1.2-fold). However, we saw a more dramatic effect of MG132 on the expression levels of unstable haplotypes I, III, and IV (3.7-, 5.6- and 4.9-fold, respectively), suggesting that these haplotypes are more actively degraded via proteasomal degradation than haplotype I (**Figure 1A**).

**Figure 1.**
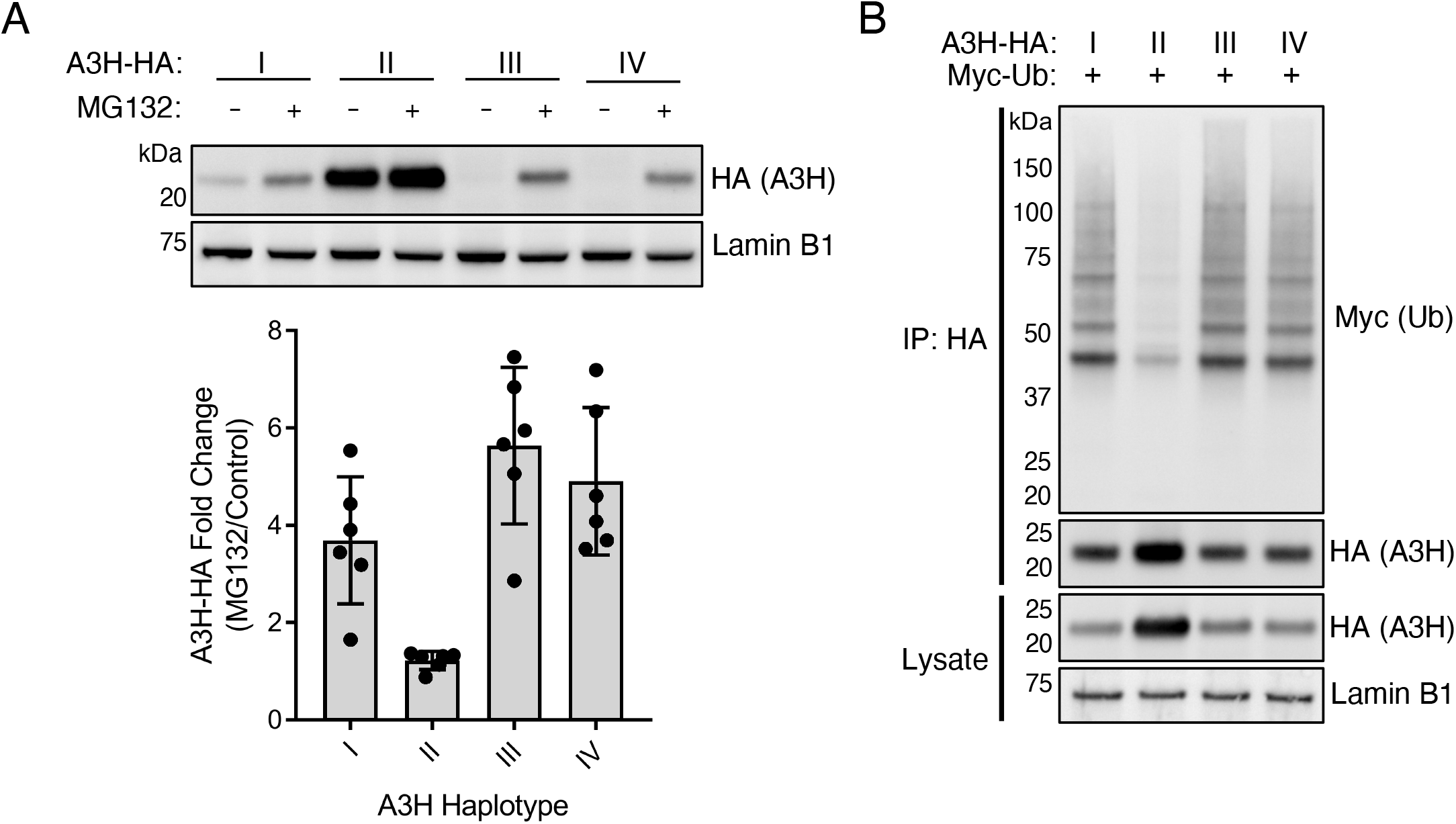
Human A3H haplotypes are differentially ubiquitinated. (**A**) Top: Immunoblot of A3H haplotypes in the absence and presence of MG132. 293T cells were transfected with plasmids expressing A3H-HA haplotypes I through IV, as indicated. Cells were then treated with MG132 (10 μM) or equal volume of DMSO for 18 h. Western blotting of whole cell lysates was performed using anti-HA to assess for A3H levels, and anti-Lamin B1 as a loading control. Bottom: Densitometric analysis of six independent experiments. Expression of A3H-HA was normalized to Lamin B1 levels and plotted as the fold change of MG132 over DMSO control. Error bars indicate standard deviation from the mean. (**B**) Immunoblot of ubiquitinated A3H haplotypes. 293T cells were co-transfected with plasmids expressing A3H-HA haplotypes, as indicated, alongside myc-tagged ubiquitin (myc-Ub). Whole cell lysates were immunoprecipitated with anti-HA resin, and western blotting was performed with anti-HA and anti-myc antibodies. Western blotting of whole cell lysates was performed using anti-HA to confirm expression of A3H, and anti-Lamin B1 as a loading control.

The increased expression of all A3H haplotypes following MG132 treatment suggests that each variant is naturally degraded to some extent through the ubiquitin proteasome system in the absence of the viral antagonist Vif. Because of the differential rates of recovery seen after MG132 treatment, we hypothesized that the rates of ubiquitination would similarly differ between protein variants. To this aim, we overexpressed each A3H-HA haplotype with myc-tagged Ubiquitin (myc-Ub). A3H-HA was immunoprecipitated and subjected to western blotting against myc-Ub. We observed higher amounts of ubiquitination for proteins encoded by haplotypes I, III and IV compared to haplotype II (**Figure 1B**: note ladder of Ub-containing proteins for haplotypes I, III, and IV). These results demonstrate that haplotypes I, III, and IV are processed by ubiquitination more heavily than haplotype II, which likely plays a large role in the turnover and loss of antiviral activity of the unstable haplotypes.

The finding that MG132 increases expression of haplotypes III and IV to levels at or exceeding that of haplotype I (**Figure 1A**) prompted us to investigate if restoring expression of N15del haplotypes would lead to an increase in antiviral activity. As MG132 has pleotropic effects and precludes reliable results from our infectivity assays, we employed the lysine mutagenesis approach previously used to study Vif-mediated polyubiquitination of A3G [32–34]. In order to prevent polyubiquitination of A3H, we converted each of the 14 lysines (K) in A3H to arginine (**Figure 2A**). By introducing arginine at these sites, we aimed to maintain the positively-charged regions thought to be important for RNA interaction and A3H dimerization [23–26]. Indeed, we found that transfection of the lysine-less mutants (K-) increased expression of proteins encoded by haplotypes III and IV (4.9-fold and 5.7-fold, respectively, **Figure 2B**). Importantly, when the lysines are mutated in any of the haplotypes ubiquitination is no longer observed (**Figure 2C**).

**Figure 2.**
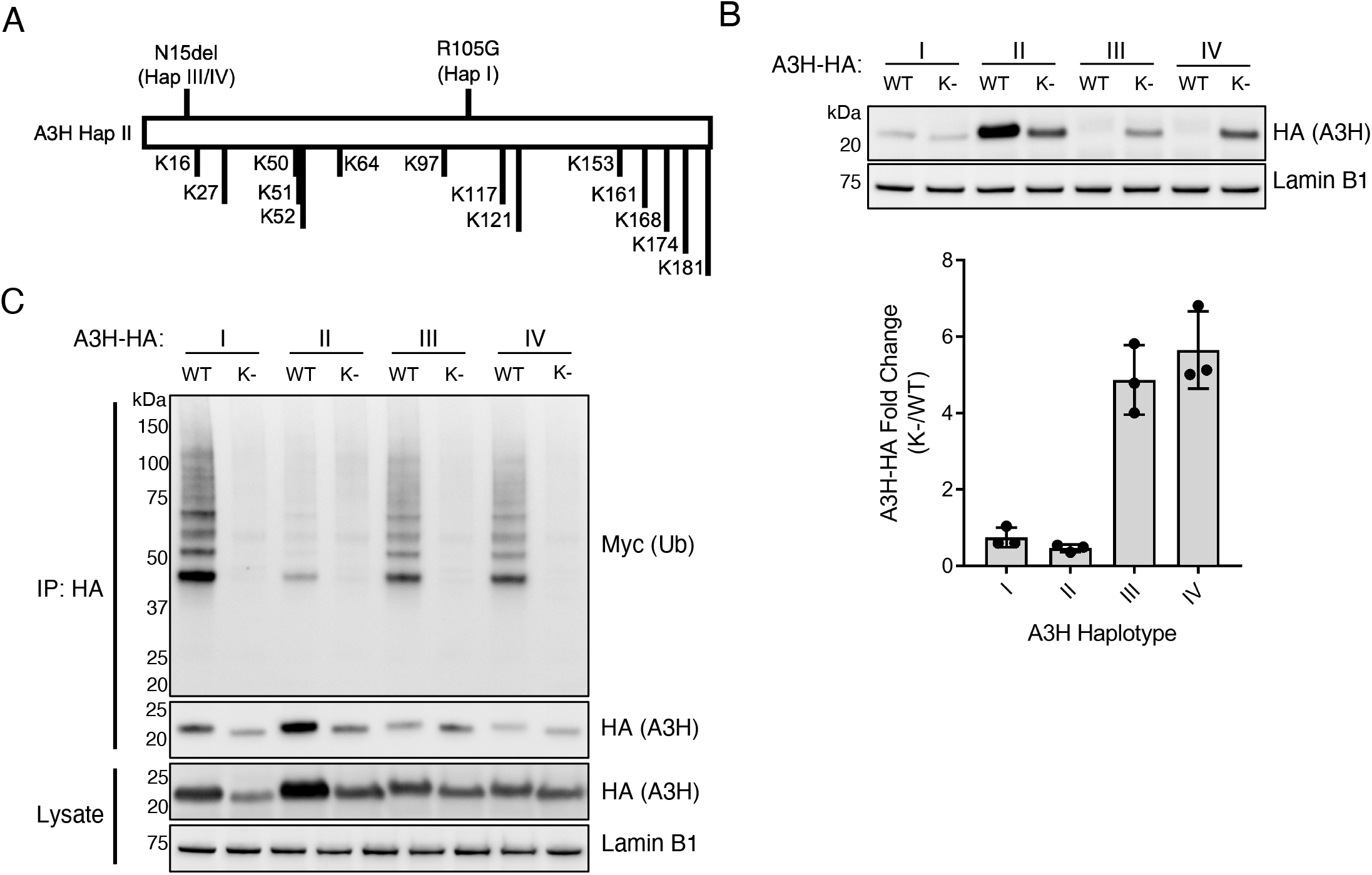
Genetic inhibition of ubiquitination increases expression of A3H haplotypes III and IV. (**A**) Protein schematic of A3H haplotype II (A3H Hap II) highlighting the polymorphic sites of haplotypes I (Hap I, R105G) and haplotypes III and IV (Hap III/IV, N15del), as well as all lysine residues (K) present in A3H. (**B**) Top: Immunoblot of wild-type (WT) and lysine-less (K-) A3H haplotypes expressed in 293T cells. Western blotting of whole cell lysates was performed using anti-HA to assess for A3H levels, and anti-Lamin B1 as a loading control. Bottom: Densitometric analysis of three independent experiments. Expression of A3H-HA was normalized to Lamin B1 levels and plotted as the fold change of lysine-less (K-) over wild-type (WT) haplotypes. Error bars indicate standard deviation from the mean. (**C**) Immunoblot of ubiquitinated wild-type (WT) and lysine-less (K-) A3H haplotypes co-transfected in 293T cells alongside myc-tagged ubiquitin (myc-Ub). Whole cell lysates were immunoprecipitated with anti-HA resin, and western blotting was performed with anti-HA and anti-myc antibodies. Western blotting of whole cell lysates was performed using anti-HA to confirm expression of A3H, and anti-Lamin B1 as a loading control.

Interestingly, while mutation of lysines increased the expression of proteins encoded by A3H haplotypes III and IV, it *decreased* the expression of proteins encoded by haplotypes I and II (**Figure 2B, 2C**). These results were unexpected, as inhibition of ubiquitin-mediated proteasomal degradation does not lead to a similar decrease in expression of these haplotypes (**Figure 1A**). It is unlikely that these mutants are toxic to cells because we did not observe increased cell death upon transfection with each haplotype and their lysine-less mutants. Another possibility is that haplotypes I and II are actively stabilized by a mechanism that requires one or more of the lysine residues, perhaps due to ubiquitination, other modifications, or structural constraints. Regardless, it is important to note that lysine mutagenesis results in two opposing phenotypes between haplotypes I and II and haplotypes III and IV.

### 3.2. Inhibiting ubiquitination does not restore antiviral activity of unstable A3H haplotypes

We next assessed if increasing the expression of unstable A3H haplotypes III and IV by preventing all ubiquitination (**Figure 2**) could rescue antiviral function. To test this, we performed single-cycle infectivity assays by expressing wild-type A3H haplotypes, or their lysine-less counterparts, alongside *vif*-deficient HIV-1. Concurrent with their significant loss of expression (**Figure 2B**), the antiviral activities of lysine-less haplotype I and haplotype II were significantly reduced compared to their wild-type counterparts (2.1-fold and 5.9-fold loss in activity, respectively) (**Figure 3A**). Importantly, in the case of haplotype II, the lysine-less mutant is still capable of reducing infection to 11.1%, suggesting that neither lysine residues nor ubiquitination are required for the majority of anti-HIV-1 activity of A3H (**Figure 3A**), and this loss of function is likely due to the decreased expression of this mutant (**Figure 2B**). However, the increased expression of haplotypes III and IV induced by mutating all lysines to arginines (**Figure 2B**), did not similarly restore anti-HIV-1 activity (**Figure 3A**). Importantly, the antiviral activities of each mutant reflect the amount of packaging into virions (**Figure 3B**). As expected, wild-type haplotype II has the highest packaging efficiency, correlating with its expression and antiviral activity, while wild-type haplotype I has far less, but detectable, expression in virions. Lysine-less mutants of haplotypes I and II each lose some degree of viral packaging, with haplotype I losing all detectable expression (**Figure 3B**). In contrast, haplotypes III and IV are barely detectable in virions with or without lysine residues (**Figure 3B**). This result suggests that the N15del mutation in haplotypes III and IV inhibits antiviral activity in a mechanism that is independent of stability, but rather is linked to a defect that prevents virion packaging.

**Figure 3.**
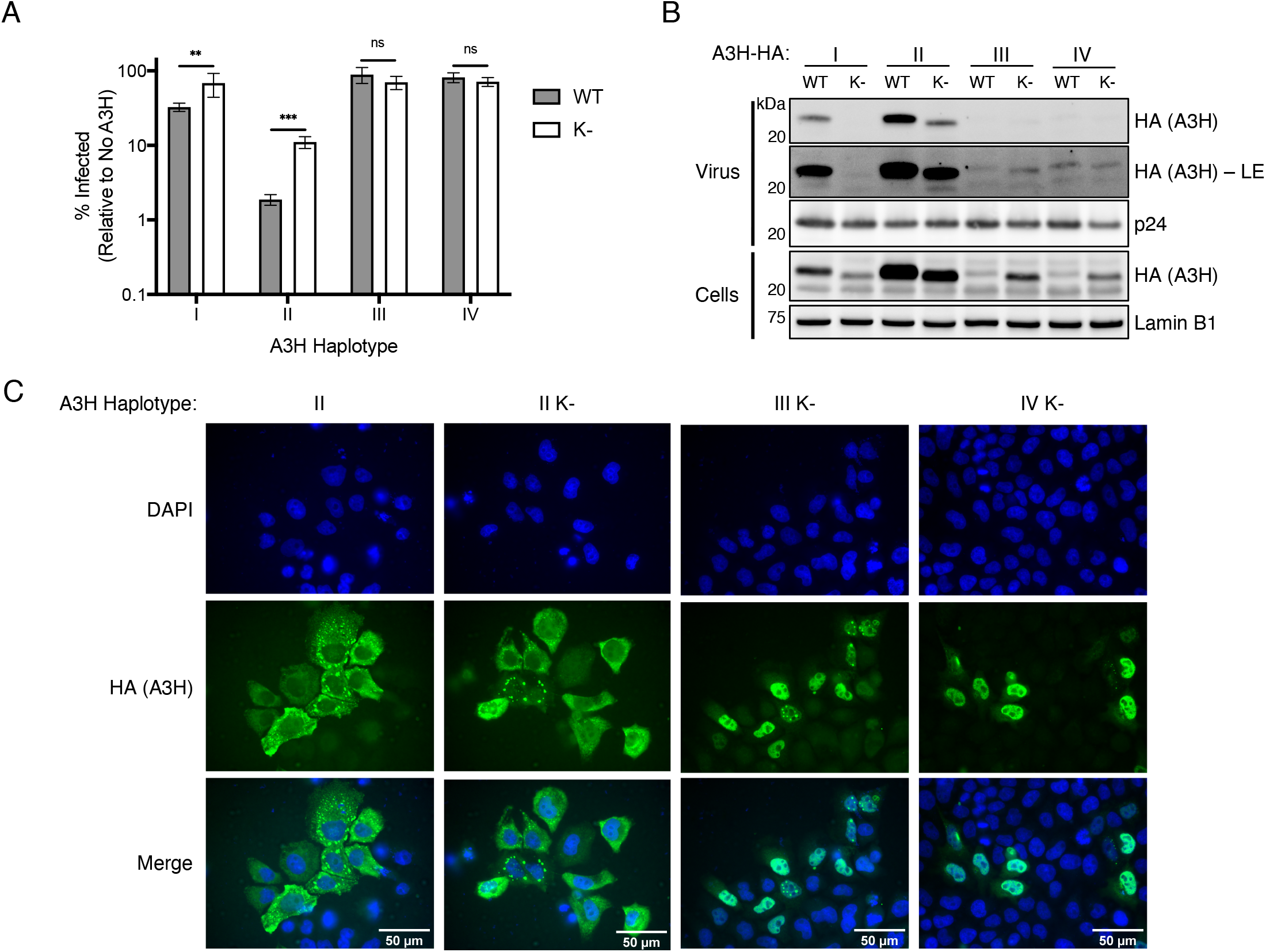
Functional deficiency of A3H haplotypes III and IV is independent of instability. (**A**) Single-cycle viral infectivity assays performed with *vif*-deficient HIV-1 virus produced in the presence of wild-type (WT, gray bars) A3H haplotypes, or lysine-less (K-, open bars) counterparts. Shown is the average of six biological replicates with error bars representing the standard deviation of the mean. **p < 0.01, ***p < 0.001, ns = not significant (p > 0.05), unpaired t test. (**B**) Immunoblot of viral supernatant (Virus, top) and whole cell lysate (Cells, bottom) of virus producing cells as in (**A**) using anti-HA to assess A3H levels packaged in virus or expressed in cells. Anti-p24 and anti-Lamin B1 were used as loading controls for virus and cell lysate, respectively. LE = long exposure. (**C**) Immunofluorescence imaging of HeLa cells transfected with wild-type A3H-HA haplotype II or lysine-less A3H-HA haplotype II (II K-), III (III K-), and IV (IV K-). Cells were stained for fluorescence microscopy using anti-HA to detect A3H (green), and DAPI to detect the nucleus (blue). Images are representative of 30 randomly selected field images across three independent experiments.

Previous studies have correlated cytoplasmic localization of A3 proteins with antiviral activity; for example, the less antiviral haplotype I has a nuclear-biased localization compared to the more cytoplasmic and active haplotype II [20]. However, due to the poor expression of haplotypes III and IV, previous studies have been unable to assess the contribution of the N15del mutation on A3H localization. By increasing expression of haplotypes III and IV through lysine mutagenesis, we reasoned we could use these mutants to determine the localization of N15del-containing variants. Importantly, the localization of lysine-less haplotype II was similar to its wild-type counterpart and was predominantly cytoplasmic, indicating that lysine mutagenesis does not alter the localization of A3H (**Figure 3C**). In contrast, immunofluorescence imaging of lysine-less haplotypes III and IV revealed a strong nuclear localization (**Figure 3C**). Taken together, these results suggest that the N15del mutation, like the R105G mutation in haplotype I [20], promotes localization of A3H to the nucleus, where it would be unable to package into nascent virions and elicit antiviral activity against HIV.

### 3.3. The stability of A3H haplotype II is dominant over the instability of haplotype III

The A3H haplotypes destabilized by the R105G (haplotype I) or N15del (haplotypes III and IV) mutations all share a nuclear-biased localization, a decrease in stability, and a decrease in antiviral activity. In a previous study, A3H constructs were made where haplotype II was linked to haplotype I, and was found that the chimeric proteins of haplotype I and II (I-II and II-I) were expressed at similar levels as haplotype II alone and much higher than haplotype I or a protein where haplotype I was duplicated (I-I) [20]. Therefore, the stability of haplotype II is dominant over the instability of haplotype I, suggesting that haplotype II is actively stabilized by a mechanism lost through R105G mutation. Taking a similar approach, we created double-domain A3H constructs to link haplotype II to haplotype III in both orientations (III-II and II-III) to address if the detrimental effects of the N15del mutation are dominant over haplotype II (**Figure 4A**).

**Figure 4.**
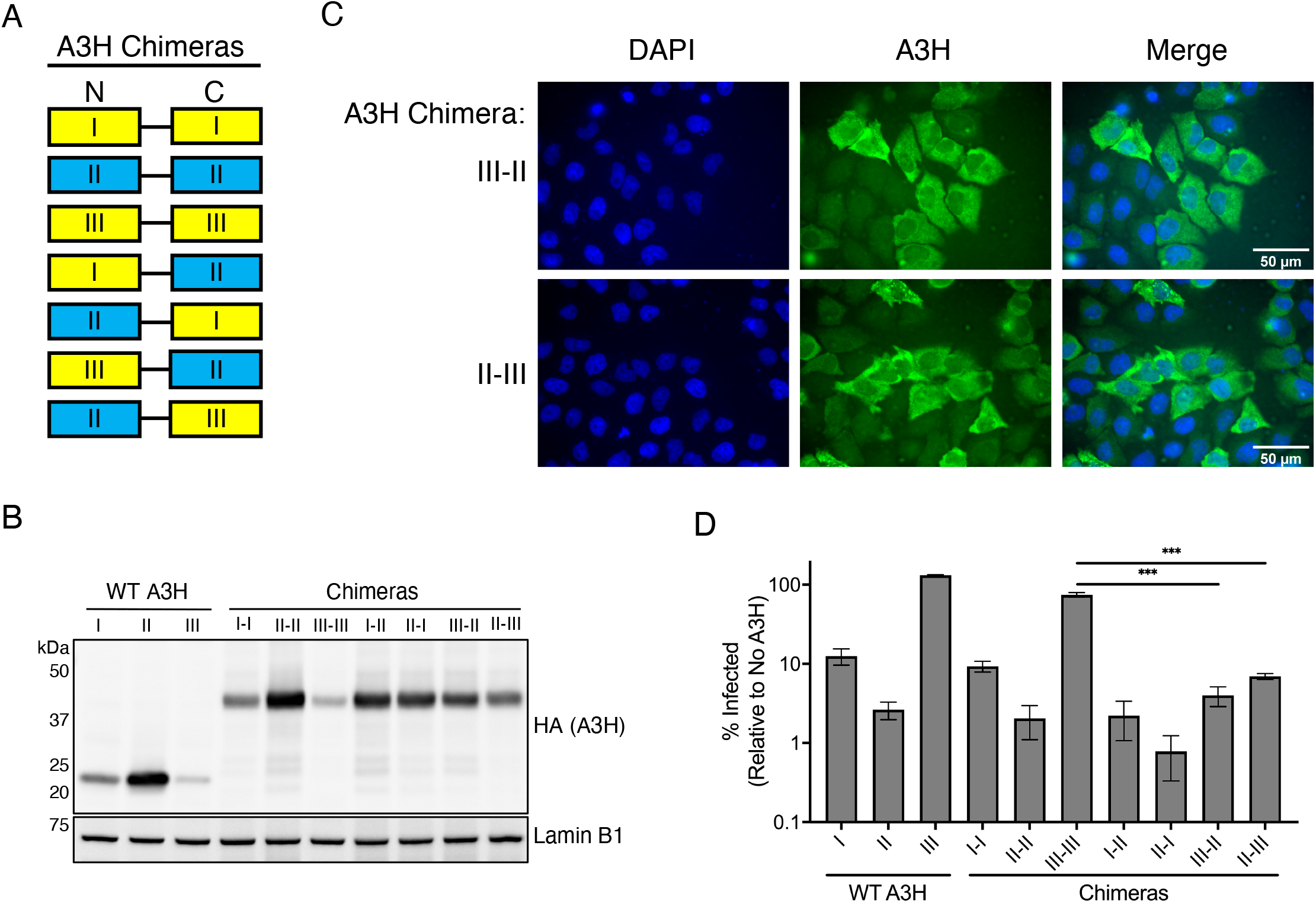
The stability and antiviral function of A3H haplotype II is dominant over the destabilizing N15del mutation of haplotype III. (**A**) Protein schematic of A3H chimeras used in this study. Haplotypes are denoted as I, II, or III in the N-terminal (N) or C-terminal (C) end of the flexible linker. Yellow boxes denote unstable haplotypes, and blue boxes represent active haplotypes. (**B**) Immunoblot of indicated wild-type (WT) A3H haplotypes or chimeras expressed in 293T cells. Western blotting of whole cell lysates was performed using anti-A3H and anti-Lamin B1 as a loading control. (**C**) Immunofluorescence imaging of HeLa cells transfected with A3H chimeras III-II, or II-III. Cells were stained for fluorescence microscopy using anti-A3H (green), and DAPI to detect the nucleus (blue). Images are representative of 30 randomly selected field images across three independent experiments. (**D**) Single-cycle infectivity assays performed with *vif*-deficient HIV-1 virus produced in the presence of the indicated wild-type A3H haplotypes or chimeras. Shown is the average of three biological replicates with error bars indicating the standard deviation of the mean. ***p < 0.001, unpaired t test.

To assess the effect of linking haplotype II to haplotype III on stability, we overexpressed these chimeras or wild-type A3H haplotypes and evaluated their expression by western blot. We anticipated one of two results: 1) the N15del mutation in haplotype III leads to active degradation even when linked to haplotype II, or 2) the active stabilization of haplotype II is dominant over the instability of haplotype III in a similar fashion to haplotype I. As shown previously, chimera I-I remains unstable, but haplotype I is stabilized (to levels similar to chimera II-II) when linked with haplotype II in either orientation (**Figure 4B**, [20]). Importantly, we note a similar increase in expression when haplotype III is linked to haplotype II in either orientation, but not when haplotype III is linked to itself, indicating that the stabilizing factors of haplotype II are also dominant over haplotype III (**Figure 4B**). Therefore, despite the more rapid turnover seen in haplotype III, association with haplotype II is sufficient to override this mechanism. The dominance of haplotype II stability over haplotype III suggests that the instability resulting from the N15del mutation is similar to that of the R105G mutation and is likely the result of passive ubiquitination and degradation rather than either of these mutations creating an active degradation signal. On the other hand, the ability of haplotype II to restore expression of both haplotypes I and III further supports the hypothesis that haplotype II stabilization is an active mechanism that correlates with protection from ubiquitination.

Given that expression of the N15del-containing haplotype III can also be increased through linkage to haplotype II (**Figure 4B**), we next performed immunofluorescence microscopy to assess the localization of these chimeras. Imaging of the III-II and II-III chimeras showed a cell-wide distribution of chimeric A3H, demonstrating that cytoplasmic retention of haplotype II is also dominant over the nuclear localization of haplotype III (**Figure 4C**). Because both the stability and cytoplasmic retention of haplotype II are dominant in these chimeras, we hypothesized that they also exhibit antiviral activity comparable to haplotype II. To this aim, we tested the ability of haplotype III chimeras to restrict *vif-*deficient HIV-1. As previously described, the antiviral activities of haplotypes I and II matched the respective activities of chimera I-I and II-II, and haplotype I linked to haplotype II (I-II and II-I) were comparably active to haplotype II (**Figure 4D** and [20]). We also found that linking haplotype III to itself (III-III), which does not rescue expression (**Figure 4B**), also does not considerably affect antiviral activity (**Figure 4D**). Finally, only linkage to haplotype II rescued antiviral activity of haplotype III against HIV-1 (**Figure 4D**). Taken together, these results indicate that the stability, cytoplasmic retention, and antiviral potency of haplotype II are dominant over both destabilizing R105G and N15del mutations. Therefore, although we find additional functional deficiencies conferred by the N15del mutation in comparison to the R105G mutation (e.g. more nuclear localization, complete loss of packaging, and no antiviral activity), these deficiencies are similarly overcome through association with haplotype II. As association with haplotype II is able to similarly overcome both instability and functional defects of R105G and N15del mutations, it is likely that the downstream effect of these mutations is due to a shared loss of active stabilization and cytoplasmic retention central to the antiviral potency of haplotype II.

## 4. Discussion

Here, we provide evidence that steady-state ubiquitination differs between the major A3H haplotypes circulating in the human population. These differences among A3H haplotypes in humans result in loss of protein expression that, only in the case of the N15del mutation, can be restored by mutation of lysines within the protein. However, the resultant protein is localized to the nucleus, fails to incorporate into virions, and does not have anti-HIV-1 activity. Our results indicate that the A3H antiviral activity was lost during human evolution, and that the inactive A3H proteins encoded by much of the human population cannot be easily restored by simply increasing the expression or by finding means to stabilize these proteins.

Ubiquitination most commonly results in proteasomal or lysosomal degradation of target proteins, although several other functions such as protein trafficking and inflammatory signaling have been described [35]. The function of ubiquitin modification on target proteins depends on the type of linkage (e.g., lysine-48 and lysine-64) and the particular lysine residues modified on the target protein. However, due to the partial recovery of unstable A3H haplotypes with the proteasomal inhibitor MG132 (**Figure 1A**), and the apparent lack of ubiquitination of the most stable A3H haplotype II (**Figure 1B**), it is likely that ubiquitination of haplotypes I, III, and IV primarily direct A3H for proteasomal degradation.

We found an intriguing phenotype by which mutating all lysines to arginine only increases expression of haplotypes III and IV, and decreases expression of I and II (**Figure 2B**). The decrease in expression observed with lysine-less haplotypes I and II was unexpected, as treatment with MG132 resulted in an increase of these two haplotypes (**Figure 1A**). Therefore, it is possible that haplotypes I and II are regulated by ubiquitin in a manner independent from proteasomal degradation or that the presence of lysine residues are otherwise important for the stability of these haplotypes.

Nonetheless, lysine-less A3H haplotype II is still potently antiviral in relation to its expression (**Figure 3A**) and maintains its cytoplasmic localization (**Figurer 3C**). However, lysine-less haplotypes III and IV express detectable proteins that allowed us to assess known functionally relevant characteristics of A3H (**Figure 2B**). Lysine-less haplotypes III and IV are almost exclusively nuclear, are unable to package into virions and have no antiviral activity (**Figure 3**). In contrast, haplotype I has some cytoplasmic localization [20,36], and low but detectable viral packaging (**Figure 3B**) and antiviral activity (**Figure 3A, 4D**). Taken together, these results reveal key differences between the R105G and N15del mutations in unstable A3H haplotypes, and suggest that the R105G mutation of haplotype I, although destabilizing, has more functional characteristics in common with the stable haplotype II than it does to the N15del-containing haplotypes III and IV.

Previous findings from our group demonstrated that the stability, cytoplasmic retention, and antiviral activity of haplotype II was dominant over the instability and nuclear bias of haplotype I by linking these haplotypes in either orientation [20]. The N15del mutation of haplotypes III and IV results in a protein that is less stable, more nuclear, and non-antiviral in comparison to haplotype I, suggesting that these haplotypes are more prone to negative regulation. Despite greater instability and functional deficiencies of N15del-containing haplotypes, we found that linking haplotype III to haplotype II in either orientation results in a stable, antiviral protein that localizes to the cytoplasm (**Figure 4**). These findings suggest that A3H variants expressing the R105G and N15del mutations may share a common negative regulatory mechanism that affects the stability and localization of these mutants, and the N15del mutation leads to greater utilization of this mechanism.

Several groups have recently described the mechanism by which stable A3H dimerizes through its interaction with duplex RNA, and this complex is maintained primarily through protein-RNA contacts [23,25–27]. Interaction with duplex RNA is mediated by aromatic and positively-charged residues in Loop 1, Loop 7, and the C-terminal α6 helix [23,25–27]. It is possible that the R105G and N15del mutations disrupt RNA binding and A3H-RNA complex formation, interfering with active stabilization and cytoplasmic retention observed with haplotype II. The R105G mutation is situated in β4 immediately upstream of the putative RNA-binding region (110-RLYYHW-115) in Loop 7 [37]. Although residue 105 is not directly implicated in RNA interaction, it is possible that mutations at R105 may alter the positioning of RNA-interacting residues in Loop 7. Importantly, the N15 residue is positioned within Loop 1, and deletion of this residue in haplotypes III and IV may directly prevent binding of A3H to its duplex RNA substrate and lead to greater instability and loss of function than what is conferred by the R105G mutation. Disrupting RNA binding prevents dimerization, and monomeric A3H may be susceptible to increased processing by ubiquitination. Indeed, mutagenesis of RNA binding residues results in decreased expression, nuclear localization, and a loss of packaging and antiviral activity [25–27], suggesting a shared mechanism of instability and loss of function to the R105G and N15del mutations. Thus, disrupting RNA binding might explain all of the phenotypes observed for the unstable and inactive haplotypes I, III, and IV.

Furthermore, the active mechanism by which haplotype II is stabilized might also be explained by RNA binding. Interestingly, five lysine residues are present in the C-terminal α6 helix in a span of 20 amino acids (**Figure 2A**), and ubiquitination of these residues may prevent duplex RNA binding. An intriguing model is that stabilization of haplotype II may result from a higher affinity of this haplotype for duplex RNA, which requires hydrogen bonding of arginine and lysine residues to RNA phosphates [23], and may preclude ubiquitination of these residues. As a result, stable and active A3H may be protected from degradative ubiquitination by nature of its unique RNA-mediated dimerization mechanism.

Interestingly, loss of A3H activity is not unique to humans, as an evolutionary analysis of A3H in Old World monkeys found that A3H activity has been lost in multiple primate species [38]. Moreover, several residues in the putative RNA binding region of Loop 1 were found to influence the antiviral potency of African green monkey and patas monkey A3H, providing further evidence that mutations introduced in this loop are a recurrent evolutionary mechanism that results in a loss of function [38]. It is unknown why these loss of function mutations of A3H arise and become fixed in multiple primate species despite the established importance of this restriction factor in the control of retroviruses. It is possible that the selective pressure to maintain A3H function has been lost due to the loss of an unknown pathogen or due to redundancy of more potent antiretroviral restriction factors such as A3G. On the other hand, functional A3H may come at a fitness cost in the absence of a pathogenic selective pressure. Although A3H haplotype I has lost much of its stability and antiviral activity, this variant has been associated with breast and lung cancer similar to the well-established cancer driver A3B [39,40]. Interestingly, a similar association has not been made for A3H haplotypes II, III, and IV, which may suggest a unique dysregulation of haplotype I promotes A3H-mediated genomic mutations driving these cancers. In contrast, the complete instability and loss of function coinciding with the N15del mutation of haplotypes III and IV may dually work to contract a redundant restriction factor while protecting the host from genomic mutations. Thus, although unstable A3H haplotypes are all ineffective against HIV, careful examination of these variants reveals differences in function that may have a significant influence on host fitness.

## Acknowledgements

We thank Jeannette Tenthorey, Molly OhAinle, and Mollie McDonnell for critical feedback of this manuscript, and members of the Emerman lab for helpful discussions. This research was supported by the NIH/NIAID (R01 AI30927 to ME), the HIV Accessory & Regulatory Complexes (HARC) Center (NIH/NIAID P50 AI150476 to ME), and a University of Washington STD/AIDS Research Training Fellowship (NIH/NIAID T32 AI07140 to NMC).

## References

1. Harris, R.S.; Dudley, J.P. APOBECs and virus restriction. Virology 2015, 479–480, 131–145.

2. Desimmie, B.A.; Delviks-Frankenberry, K.A.; Burdick, R.; Qi, D.; Izumi, T.; Pathak, V.K. Multiple APOBEC3 Restriction Factors for HIV-1 and One Vif to Rule Them All. J. Mol. Biol. 2014, 426, 1220–1245.

3. Sawyer, S.L.; Emerman, M.; Malik, H.S. Ancient Adaptive Evolution of the Primate Antiviral DNA-Editing Enzyme APOBEC3G. PLoS Biol. 2004, 2.

4. McLaughlin, R.N.; Gable, J.T.; Wittkopp, C.J.; Emerman, M.; Malik, H.S. Conservation and Innovation of APOBEC3A Restriction Functions during Primate Evolution. Mol. Biol. Evol. 2016, 33, 1889–1901.

5. Nakano, Y.; Aso, H.; Soper, A.; Yamada, E.; Moriwaki, M.; Juarez-Fernandez, G.; Koyanagi, Y.; Sato, K. A conflict of interest: the evolutionary arms race between mammalian APOBEC3 and lentiviral Vif. Retrovirology 2017, 14, 31.

6. Feng, Y.; Baig, T.T.; Love, R.P.; Chelico, L. Suppression of APOBEC3-mediated restriction of HIV-1 by Vif. Front. Microbiol. 2014, 5.

7. Shandilya, S.M.D.; Bohn, M.-F.; Schiffer, C.A. A computational analysis of the structural determinants of APOBEC3’s catalytic activity and vulnerability to HIV-1 Vif. Virology 2014, 471–473, 105–116.

8. Jäger, S.; Kim, D.Y.; Hultquist, J.F.; Shindo, K.; LaRue, R.S.; Kwon, E.; Li, M.; Anderson, B.D.; Yen, L.; Stanley, D.; et al. Vif hijacks CBF-β to degrade APOBEC3G and promote HIV-1 infection. Nature 2012, 481, 371–375.

9. Kim, D.Y.; Kwon, E.; Hartley, P.D.; Crosby, D.C.; Mann, S.; Krogan, N.J.; Gross, J.D. CBFβ Stabilizes HIV Vif to Counteract APOBEC3 at the Expense of RUNX1 Target Gene Expression. Mol. Cell 2013, 49, 632–644.

10. Yu, X.; Yu, Y.; Liu, B.; Luo, K.; Kong, W.; Mao, P.; Yu, X.-F. Induction of APOBEC3G ubiquitination and degradation by an HIV-1 Vif-Cul5-SCF complex. Science 2003, 302, 1056–1060.

11. Zhang, W.; Du, J.; Evans, S.L.; Yu, Y.; Yu, X.-F. T-cell differentiation factor CBF-β regulates HIV-1 Vif-mediated evasion of host restriction. Nature 2012, 481, 376–379.

12. Compton, A.A.; Hirsch, V.M.; Emerman, M. The Host Restriction Factor APOBEC3G and Retroviral Vif Protein Coevolve due to Ongoing Genetic Conflict. Cell Host Microbe 2012, 11, 91–98.

13. Compton, A.A.; Emerman, M. Convergence and Divergence in the Evolution of the APOBEC3G-Vif Interaction Reveal Ancient Origins of Simian Immunodeficiency Viruses. PLOS Pathog. 2013, 9, e1003135.

14. Ebrahimi, D.; Richards, C.M.; Carpenter, M.A.; Wang, J.; Ikeda, T.; Becker, J.T.; Cheng, A.Z.; McCann, J.L.; Shaban, N.M.; Salamango, D.J.; et al. Genetic and mechanistic basis for APOBEC3H alternative splicing, retrovirus restriction, and counteraction by HIV-1 protease. Nat. Commun. 2018, 9, 1–11.

15. Harari, A.; Ooms, M.; Mulder, L.C.F.; Simon, V. Polymorphisms and Splice Variants Influence the Antiretroviral Activity of Human APOBEC3H. J. Virol. 2009, 83, 295–303.

16. OhAinle, M.; Kerns, J.A.; Li, M.M.H.; Malik, H.S.; Emerman, M. Anti-retroelement Activity of APOBEC3H was Lost Twice in Recent Human Evolution. Cell Host Microbe 2008, 4, 249–259.

17. Refsland, E.W.; Hultquist, J.F.; Luengas, E.M.; Ikeda, T.; Shaban, N.M.; Law, E.K.; Brown, W.L.; Reilly, C.; Emerman, M.; Harris, R.S. Natural Polymorphisms in Human APOBEC3H and HIV-1 Vif Combine in Primary T Lymphocytes to Affect Viral G-to-A Mutation Levels and Infectivity. PLOS Genet. 2014, 10, e1004761.

18. Wang, X.; Abudu, A.; Son, S.; Dang, Y.; Venta, P.J.; Zheng, Y.-H. Analysis of Human APOBEC3H Haplotypes and Anti-Human Immunodeficiency Virus Type 1 Activity. J. Virol. 2011, 85, 3142–3152.

19. Zhang, Z.; Gu, Q.; Montero, M. de M.; Bravo, I.G.; Marques-Bonet, T.; Häussinger, D.; Münk, C. Stably expressed APOBEC3H forms a barrier for cross-species transmission of simian immunodeficiency virus of chimpanzee to humans. PLOS Pathog. 2017, 13, e1006746.

20. Li, M.M.H.; Emerman, M. Polymorphism in human APOBEC3H affects a phenotype dominant for subcellular localization and antiviral activity. J. Virol. 2011, 85, 8197–8207.

21. Tan, L.; Sarkis, P.T.N.; Wang, T.; Tian, C.; Yu, X.-F. Sole copy of Z2-type human cytidine deaminase APOBEC3H has inhibitory activity against retrotransposons and HIV-1. FASEB J. 2008, 23, 279–287.

22. OhAinle, M.; Kerns, J.A.; Malik, H.S.; Emerman, M. Adaptive Evolution and Antiviral Activity of the Conserved Mammalian Cytidine Deaminase APOBEC3H. J. Virol. 2006, 80, 3853–3862.

23. Bohn, J.A.; Thummar, K.; York, A.; Raymond, A.; Brown, W.C.; Bieniasz, P.D.; Hatziioannou, T.; Smith, J.L. APOBEC3H structure reveals an unusual mechanism of interaction with duplex RNA. Nat. Commun. 2017, 8, 1–9.

24. Ito, F.; Yang, H.; Xiao, X.; Li, S.-X.; Wolfe, A.; Zirkle, B.; Arutiunian, V.; Chen, X.S. Understanding the Structure, Multimerization, Subcellular Localization and mC Selectivity of a Genomic Mutator and Anti-HIV Factor APOBEC3H. Sci. Rep. 2018, 8.

25. Shaban, N.M.; Shi, K.; Lauer, K.V.; Carpenter, M.A.; Richards, C.M.; Salamango, D.; Wang, J.; Lopresti, M.W.; Banerjee, S.; Levin-Klein, R.; et al. The Antiviral and Cancer Genomic DNA Deaminase APOBEC3H Is Regulated by an RNA-Mediated Dimerization Mechanism. Mol. Cell 2018, 69, 75–86.e9.

26. Matsuoka, T.; Nagae, T.; Ode, H.; Awazu, H.; Kurosawa, T.; Hamano, A.; Matsuoka, K.; Hachiya, A.; Imahashi, M.; Yokomaku, Y.; et al. Structural basis of chimpanzee APOBEC3H dimerization stabilized by double-stranded RNA. Nucleic Acids Res. 2018, 46, 10368–10379.

27. Feng, Y.; Wong, L.; Morse, M.; Rouzina, I.; Williams, M.C.; Chelico, L. RNA-Mediated Dimerization of the Human Deoxycytidine Deaminase APOBEC3H Influences Enzyme Activity and Interaction with Nucleic Acids. J. Mol. Biol. 2018, 430, 4891–4907.

28. Li, M.M.H.; Wu, L.I.; Emerman, M. The Range of Human APOBEC3H Sensitivity to Lentiviral Vif Proteins. J. Virol. 2010, 84, 88–95.

29. Vermeire, J.; Naessens, E.; Vanderstraeten, H.; Landi, A.; Iannucci, V.; Nuffel, A.V.; Taghon, T.; Pizzato, M.; Verhasselt, B. Quantification of Reverse Transcriptase Activity by Real-Time PCR as a Fast and Accurate Method for Titration of HIV, Lenti- and Retroviral Vectors. PLOS ONE 2012, 7, e50859.

30. Schneider, C.A.; Rasband, W.S.; Eliceiri, K.W. NIH Image to ImageJ: 25 years of image analysis. Nat. Methods 2012, 9, 671–675.

31. Dang, Y.; Siew, L.M.; Wang, X.; Han, Y.; Lampen, R.; Zheng, Y.-H. Human cytidine deaminase APOBEC3H restricts HIV-1 replication. J. Biol. Chem. 2008, 283, 11606–11614.

32. Dang, Y.; Siew, L.M.; Zheng, Y.-H. APOBEC3G Is Degraded by the Proteasomal Pathway in a Vif-dependent Manner without Being Polyubiquitylated. J. Biol. Chem. 2008, 283, 13124–13131.

33. Turner, T.; Shao, Q.; Wang, W.; Wang, Y.; Wang, C.; Kinlock, B.; Liu, B. Differential contributions of ubiquitin-modified APOBEC3G lysine residues to HIV-1 Vif-induced degradation. J. Mol. Biol. 2016, 428, 3529–3539.

34. Wang, Y.; Shao, Q.; Yu, X.; Kong, W.; Hildreth, J.E.K.; Liu, B. N-Terminal Hemagglutinin Tag Renders Lysine-Deficient APOBEC3G Resistant to HIV-1 Vif-Induced Degradation by Reduced Polyubiquitination. J. Virol. 2011, 85, 4510–4519.

35. Swatek, K.N.; Komander, D. Ubiquitin modifications. Cell Res. 2016, 26, 399–422.

36. Ooms, M.; Majdak, S.; Seibert, C.W.; Harari, A.; Simon, V. The Localization of APOBEC3H Variants in HIV-1 Virions Determines Their Antiviral Activity. J. Virol. 2010, 84, 7961–7969.

37. Mitra, M.; Singer, D.; Mano, Y.; Hritz, J.; Nam, G.; Gorelick, R.J.; Byeon, I.-J.L.; Gronenborn, A.M.; Iwatani, Y.; Levin, J.G. Sequence and structural determinants of human APOBEC3H deaminase and anti-HIV-1 activities. Retrovirology 2015, 12, 3.

38. Garcia, E.I.; Emerman, M. Recurrent Loss of APOBEC3H Activity during Primate Evolution. J. Virol. 2018, 92.

39. Burns, M.B.; Lackey, L.; Carpenter, M.A.; Rathore, A.; Land, A.M.; Leonard, B.; Refsland, E.W.; Kotandeniya, D.; Tretyakova, N.; Nikas, J.B.; et al. APOBEC3B is an enzymatic source of mutation in breast cancer. Nature 2013, 494, 366–370.

40. Starrett, G.J.; Luengas, E.M.; McCann, J.L.; Ebrahimi, D.; Temiz, N.A.; Love, R.P.; Feng, Y.; Adolph, M.B.; Chelico, L.; Law, E.K.; et al. The DNA cytosine deaminase APOBEC3H haplotype I likely contributes to breast and lung cancer mutagenesis. Nat. Commun. 2016, 7, 1–13.

